# Bias in community-weighted mean analysis of plant functional traits and species indicator values

**DOI:** 10.1101/046946

**Authors:** David Zelený

## Abstract

One way to analyze the relationship between species attributes and sample attributes via the matrix of species composition is to calculate the community-weighted mean of species attributes (CWM) and relate it to sample attributes by correlation, regression or ANOVA. This *weighted-mean approach* is frequently used by vegetation ecologists to relate species attributes like plant functional traits or Ellenberg-like species indicator values to sample attributes like measured environmental variables, biotic properties, species richness or sample scores in ordination analysis.

The problem with the weighted-mean approach is that, in certain cases, it yields biased results in terms of both effect size and *P*-values, and this bias is contingent upon the beta diversity of the species composition data. The reason is that CWM values calculated from samples of communities sharing some species are not independent of each other. This influences the number of effective degrees of freedom, which is usually lower than the actual number of samples, and the difference further increases with decreasing beta diversity of the data set. The discrepancy between the number of effective degrees of freedom and the number of samples in analysis turns into biased effect sizes and an inflated Type I error rate in those cases where the significance of the relationship is tested by standard tests, a problem which is analogous to analysis of two spatially autocorrelated variables. Consequently, results of studies using rather homogeneous (although not necessarily small) compositional data sets may be overly optimistic, and effect sizes of studies based on data sets differing by their beta diversity are not directly comparable.

Here, I introduce guidelines on how to decide in which situation the bias is actually a problem when interpreting results, recognizing that there are several types of species and sample attributes with different properties and that ecological hypotheses commonly tested by the weighted-mean approach fall into one of three broad categories. I also compare available analytical solutions accounting for the bias (modified permutation test and sequential permutation test using the fourth-corner statistic) and suggest rules for their use.

Abbreviations

CWM –: community-weighted mean.

## Introduction

A common task of community ecologists is to relate species attributes (like species functional traits) to sample attributes (such as environmental variables) using data about species composition of local community samples. One way to do it is by calculating the community-weighted mean (CWM) of species attributes for each sample, relate it directly to sample attributes by correlation, regression or ANOVA, and test the significance of this relationship. Although frequently used, this *weighted-mean approach* has some serious limitations, and researchers using it should be aware of them. Most notably, in situations discussed here in detail, the results appear more optimistic than they in reality are, i.e. they are biased in terms of estimated effect size and Type I error rate.

Generally, in weighted-mean approach, *species attributes* could be species properties (traits), behavior (species ecological optima) or phylogenetic age, while *sample attributes* are measured or estimated characteristics of community samples (environmental variables) or variables derived from a matrix of species composition (like species richness or positions of samples in ordination diagrams). In vegetation ecology, the two types of broadly used species attributes are plant functional traits and species indicator values. CWMs of plant functional traits can be related to environmental variables to demonstrate the effect of environmental filtering on trait-mediated community assembly (Díaz et al. 1998; Shipley 2010), or to predict changes in ecosystem properties, such as biomass production or nutrient cycling (Garnier et al. 2004; Vile et al. 2006), or ecosystem services like fodder production or maintenance of soil fertility (Díaz et al. 2007). CWMs of species indicator values (e.g. those of Ellenberg et al. 1992) are used to estimate habitat conditions from known species composition of vegetation samples, and these estimates are often related to soil, light or climatic variables (Schaffers & Sýkora 2000) or used for ecological interpretation of compositional changes (e.g. by relating them to axes of unconstrained ordination, Persson 1981). Other, more specific examples include relating the community specialization index (CSI, mean of species specialization values weighted by their dominance in the community) to environmental variables (Clavero & Brotons 2010; Fajmonová et al. 2013; Carboni et al. 2016), or attempts to verify whether plant biomass can be estimated from tabulated plant heights and species composition as the mean of species heights weighted by their cover in a plot (Axmanová et al. 2012).

Weighted-mean approach is also used in other fields, like biogeography (relating grid-based means of species properties, such as animal body size, to macroclimate or diversity; Hawkins & Diniz-Filho 2006), hydrobiology (relating trophic diatom index based on weighted mean of diatom indicator values to measured water quality parameters to assess its reliability; Kelly & Whitton 1995), or paleoecology (one of the transfer functions used to reconstruct acidification of lakes from fossil diatom assemblages preserved in lake sediments is based on weighted means of diatom optima along the pH gradient; ter Braak & Barendregt 1986; Birks et al. 1990).

Although the weighted-mean approach technically relates two sets of variables (CWM and sample attributes), three matrices are in fact involved in the computation background (notation here follows the RLQ analysis of Dolédec et al. 1996): matrix of *sample attributes* **R** with *m* sample attributes of *n* samples (*n* × *m*); matrix of *species composition* **L** with abundance (or presence-absence) of *p* species in *n* samples (*n* × *p*); and matrix of *species attributes* **Q** with *s* species attributes for *p* species (*s* × *p*). The weighted-mean approach is just one of the possible options for relating species attributes (**Q**) to sample attributes (**R**) via a matrix of species composition (**L**): it combines **Q** with **L** into a matrix of weighted-means (**M**) and relates it to **R**. An alternative solution is to combine a matrix of sample attributes **R** with species composition **L** by calculating the weighted-mean of sample attributes (optima of individual species along a given sample attribute or species centroids) and relate these values to species attributes **Q** (e.g. ter Braak & Looman 1986). The third option is to use methods suitable for simultaneously handling all three matrices (**R**, **L** and **Q**), such as the *fourth-corner approach* (Legendre et al. 1997), the related ordination method, called RLQ analysis (Dolédec et al. 1996), or other alternatives (Jamil et al. 2013, Brown et al. 2014).

In the weighted-mean approach, the relationship between CWM and sample attributes, after being analyzed by correlation, regression or ANOVA, is often tested by a standard parametric or permutation test (called simply *standard test* throughout this study). However, as will be demonstrated further in this paper, not all types of ecological questions analyzed by the weighted-mean approach should be tested by the standard test, since it may generate biased results. This problem was pointed out by Jansen et al. (2011) and Zelený & Schaffers (2012) in the context of mean Ellenberg indicator values, by Peres-Neto et al. (2012, 2016) in the context of metacommunity phylogenetics and species functional traits, by Šmilauer & Lepš (2014, p. 158) in the context of the CWM-RDA method (Nygaard & Ejrnaes 2004), and by Hawkins et al. (2017) in the macroecological context. Zelený & Schaffers (2012) suggested to solve the bias by *modified permutation test*, an alternative to the standard permutation test between CWM and sample attributes, in which species attributes instead of sample attributes are permuted. Peres-Neto et al. (2016) introduced *sequential test* (ter Braak et al. 2012), using the *fourth-corner* statistic (Legendre et al. 1997).

Here, I justify the source of the bias, among others also using an analogy with the bias in the analysis of spatially autocorrelated data, and clarify when the bias is an issue and when it is not. For this, I define several types of species and sample attributes, differing by their origin and relationship to a matrix of species composition. Then I review which ecological questions and null hypotheses are commonly analyzed by the weighted-mean approach and sort them into three broad categories. Using simulated data, I show for which of these categories there is a risk of biased results if tested by standard test and how this bias depends on the beta diversity of species composition matrix. Finally, I review and compare methods available for solving the problem of inflated Type I error rate in the weighted-mean approach, namely the *modified permutation test* (Zelený & Schaffers 2012) and the *sequential permutation test* based on the *fourth-corner statistic* (Peres-Neto et al. 2016), and suggest guidelines for their use. Although all examples and reasoning used here are focused on the relationship of species functional traits or Ellenberg-like species indicator values with sample attributes analyzed by the weighted-mean approach, the general concept is also valid for other types of species and sample attributes linked by the weighted-mean approach.

## Justification of the bias

Since the CWM of species attributes are calculated from species attributes assigned to individual species and from species composition of individual community samples, they inherit information from both sources. The numerical difference between CWM values calculated from two community samples is necessarily constrained by a difference in their species composition. Two samples with identical species composition (or, more precisely, identical relative species abundances) have identical calculated weighted-means, and two samples with slightly different species composition have CWM values rather similar. Therefore, two CWM values are not independent from each other if they are calculated from samples that share some species, and they do not bring two independent degrees of freedom into the analysis. This is because the CWM value of one sample is to some extent predictable from the CWM of the other sample from known differences in their species composition. If sample attributes are also in some predictable way related to species composition and therefore not independent from each other, their analysis with CWM becomes problematic. Since for standard parametric test the number of degrees of freedom is important for choosing the correct statistical distribution for a given sample size, disparity between the real number and effective number of samples (and degrees of freedom) leads to the selection of narrower confidence intervals and hence a higher probability of obtaining significant results (Legendre & Legendre 2012). This problem scales up to the data set level: in the case of two compositional data sets with the same number of samples used in the weighted-mean approach, the one with lower beta diversity (with samples sharing more species) has a lower number of effective degrees of freedom compared to the one with higher beta diversity.

The source of bias in weighted-mean approach can also be understood from the analogy with analysis of the relationship between two spatially autocorrelated variables. For each spatially autocorrelated variable, samples located nearby in geographical space have more similar values than expected if the values are randomly selected, and therefore they are not statistically independent (Legendre 1993). A new observation does not bring entirely new information, because its value can be partly derived from the value observed at a nearby site, and the effective number of samples (and the effective number of degrees of freedom) is lower than the real number of samples. If both variables are spatially autocorrelated, analysis of their relationship will yield biased results (unreliable effect size and inflated Type I error rate; Legendre 1993). If only one or none of the variable are spatially autocorrelated, the bias caused by autocorrelation does not appear. In the case of the weighted-mean approach, it is not the proximity in a geographical space, but the proximity in a compositional space, which reflects compositional similarities between pairs of samples. Compositional space can be imagined as an ordination diagram in which distances between plots reflect their composition dissimilarity. The bias is present only if *both variables* (CWM and sample attributes) are autocorrelated in the compositional space, i.e. if CWM values are calculated from species composition data in which some samples share some species, and sample attributes are (in some predictable way) related to species composition. If either CWM values or sample attributes (or both) are not linked to species composition, the issue with a bias does not apply. For CWM values, this can happen in the (rather unlikely) case that individual samples in the species composition matrix do not share any species and calculated CWM values are therefore completely independent from each other. For sample attributes, the question how to recognize whether they are (or should be regarded as) linked to species composition will be elaborated in the next section.

## Types of species and sample attributes

When considering alternative types of questions commonly analyzed by the weighted-mean approach, it proves useful to classify species and sample attributes according to their relationship to the matrix of species composition (**L**) as *internal* or *external*, and *linked* or *not linked* (to species composition). While the *internal/external* distinction refers to the origin of attributes (whether they are numerically derived from the matrix of species composition or not), the *linked/not linked* distinction depends on an assumed link of these attributes to the matrix of species composition.

*Internal* attributes are numerically (or in another deterministic way) derived from the matrix of species composition, while *external* attributes are typically measured or estimated variables obtained independently on the matrix of species composition. *Internal species* attributes are, e.g., species optima calculated as the weighted means of sample attributes or as species scores on ordination axes, and similarly *internal sample* attributes are sample scores on ordination axes, species richness of individual samples or the assignment of samples into groups based on compositional similarity (e.g. by numerical classification or by expert assignment based on actual species composition). *External species* attributes, on the other hand, are measured traits or tabulated species indicator values, and *external sample* attributes are measured or estimated environmental variables or assignment of samples to experimental treatments.

The distinction between attributes *linked* or *not linked* to the matrix of species composition is more subtle and depends on the context of the study. The link of (species or sample) attributes to the matrix of species composition can be acknowledged explicitly by the hypothesis we aim to test (e.g. we assume that studied traits are functional, which is why we consider them as linked to the matrix of species composition), or implicitly by the context of the study (e.g. from the experimental design). To help with the decision whether attributes are linked to species composition or not, we may ask whether it would be relevant to randomize given attributes for the purpose of testing the null hypothesis; such randomization breaks the link of (species or sample) attributes to species composition and allows to test whether the real (not randomized) attributes are linked to species composition. Those attributes, for which randomization is not relevant, should be considered as linked to species composition. From this logic, *internal* attributes should always be considered as linked to species composition, because it would not be relevant to randomize them (the null hypothesis “attributes derived from species composition are not linked to species composition” would be easy to reject). Attributes *not linked* to species composition are those which are not acknowledged (explicitly or implicitly) by the tested hypothesis or context of the study, and whose randomization would be relevant. *Linked species attributes* are, for example, species traits which are directly measured on individuals of given dataset and are considered as functional, i.e. directly influencing species performance. *Linked sample attributes* are assignments of plots into categories of experimental design (e.g. in fertilized experiment) in case that we are not questioning the effect of treatments on species composition, but instead asking how is this effect reflected by changes in species attributes (e.g. by Ellenberg-like species indicator values or traits). Examples in categories of internal species and sample attributes listed in the previous paragraphs are also examples of linked species and sample attributes, respectively. An example of *not-linked species attributes* are Ellenberg-like indicator values, which usually originate from a different context (independent from analyzed compositional data set) and even different regions. *Not-linked sample attributes* are various measured or estimated environmental factors and variables derived from GIS layers.

Sometimes the decision whether given attribute is external or internal and linked or not linked to species composition may be rather subjective. For example, an assignment of samples into habitat or vegetation types could be considered as external sample attribute (not linked to species composition). However, if this assignment is based on actual species composition of samples (or even derived from numerical classification of species composition matrix), it would be more relevant to consider them as internal sample attributes (and hence linked to species composition). Similarly, species traits, which are compiled from extensive trait databases for which we are unsure whether they are functional or not, should in most cases be considered as not linked to species composition, since there is not enough justification to think otherwise.

## Three categories of hypotheses tested by weighted-mean approach

Considering the distinction between linked and not-linked (sample or species) attributes, hypotheses commonly tested by the weighted-mean approach fall into one of the three categories (see Table 1 for a summary). *Category A* assumes that species attributes are not linked, and sample attributes are linked to the matrix of species composition. The assumption of *category B* is opposite, with species attributes linked, and sample attributes not linked. Finally, *category C* does not assume any link of either species or sample attributes to the matrix of species composition. Below, I review assumptions of ecological questions in individual categories, formulate the null hypotheses which are being tested, and suggest some examples for each of them.

**Table 1.** Overview of the characteristics of the three categories of hypotheses tested by the *weighted-mean* approach. For each category, the corresponding assumption about a link between sample attributes (**R**) or species attributes (**Q**) and species composition (**L**) is provided, as well as the null vs alternative hypothesis, a scenario within the simulated data relevant in the context of a given category (see Fig. 1), and the recommended test (standard: standard parametric or permutation test; modified: modified permutation test; sequential with 4c: the sequential permutation test with the fourth-corner statistic).

| Category of hypotheses |  | A | B | C |
| --- | --- | --- | --- | --- |
| **Assumption** |  | sample attributes linked to the matrix of species composition (**R** <--> **L**) | species attributes linked to the matrix of species composition (**Q** <--> **L**) | no assumption about the link of species or sample attributes to the matrix of species composition |
| **Null hypothesis** |  | **Q** <-//-> **L** | **R** <-//-> **L** | **R** <-//-> **Q**,<br>i.e. **R** <-//-> **L** and/or **Q** <-//-> **L** |
| **Alternative hypothesis** |  | **Q** <--> **L** | **R** <--> **L** | **R** <--> **Q**,<br>i.e. **R** <--> **L** and **Q** <--> **L** |
| **Relevant scenario(s)** |  | Scenario 2 | Scenario 3 | Scenarios 2, 3 and 4 |
| **Recommended test** | **standard** | no (biased result) | yes | no (biased result) |
|  | **modified** | yes | no | yes* |
|  | **sequential with 4c** | yes | yes | yes* |

\* too conservative if the beta diversity of the species composition matrix is low
<-//-> - no link between the two matrices, <--> - link between the two matrices.

**Figure 1.**
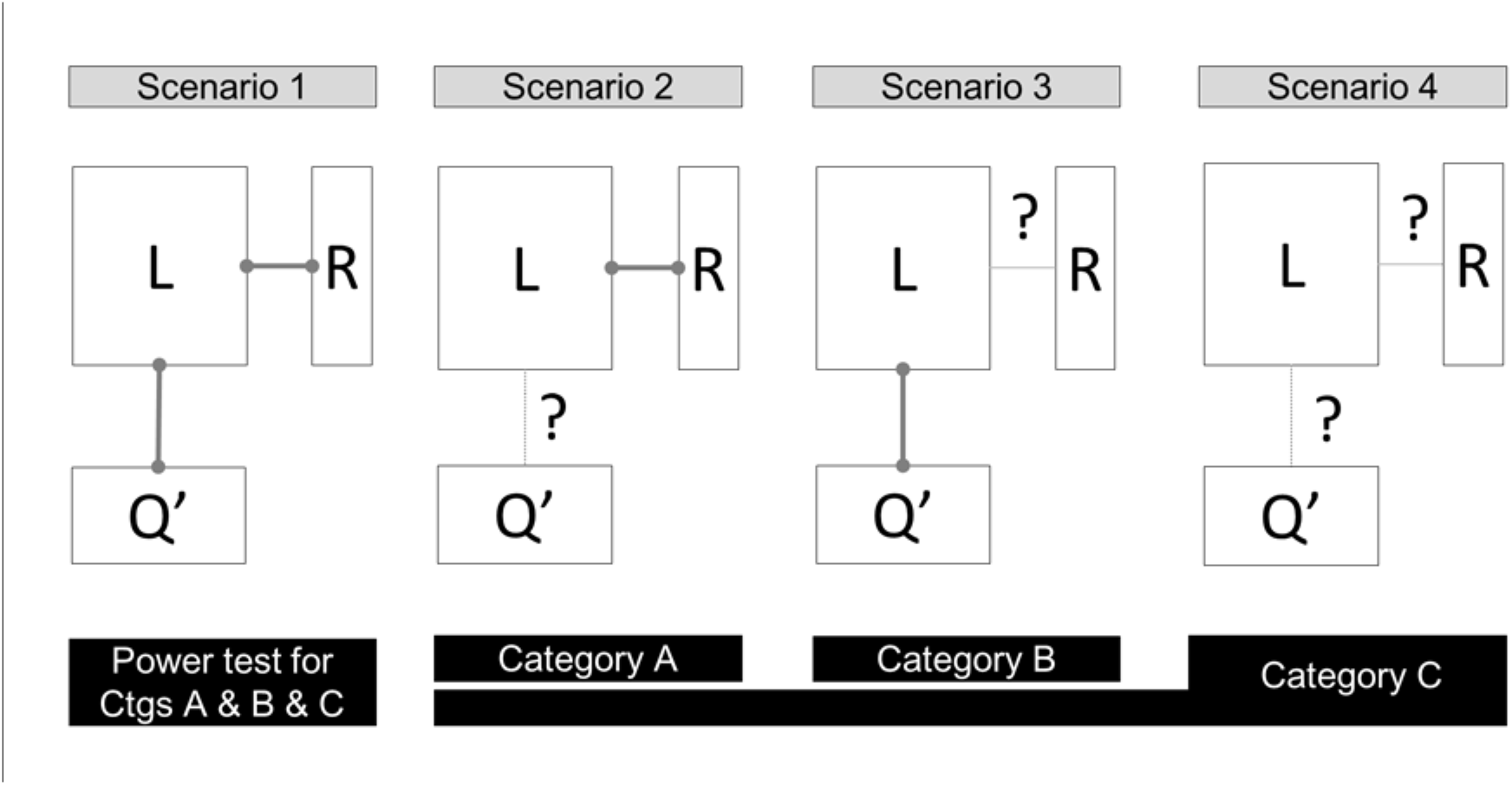
Conceptual differences between scenarios 1 to 4 in the simulated community data and the link of these scenarios (gray boxes at the top) to categories A to C (black boxes at the bottom). In scenario 1, both sample attributes (**R**) and species attributes (**Q**) are linked to the matrix of species composition (**L**), while in the other three scenarios one or both types of attributes are without the link to species composition (notified by “?” in the schema). The link of attributes to species composition was removed by permuting the values of species attributes (scenario 2), sample attributes (scenario 3) or both (scenario 4). The matrix of species attributes is transposed (**Q’**) to match the dimension of the matrix of species composition (**L**). In simulated data, both sample and species attributes are represented by a matrix of only one column (yet still using the notation for a matrix).

### Category A – sample attributes linked to species composition

Ecological questions in this category explicitly acknowledge the link between sample attributes and species composition, or the link is implicit from the context or the numerical background of the study. What is tested here is the link of species attributes to species composition. The null hypothesis states that species attributes are not linked to species composition, while the alternative hypothesis states that they are. This category includes studies focused on relating CWM to internal sample attributes, i.e. those derived numerically from the matrix of species composition (and therefore linked to it). Examples include relating mean Ellenberg indicator values to sample scores in unconstrained ordination to interpret the ecological meaning of ordination axes (Zelený & Schaffers 2012) or relating mean trait values to species richness (Hawkins et al. 2017). Studies with external sample attributes fall into this category if the sample attributes are considered linked to species composition, as is the case of experimental study in which the effect of experimental treatment (sample attribute) is acknowledged, and the question is about the response of CWM to it. An example includes the test how mean Ellenberg values reflect the changes in grassland species composition following experimental fertilizer application (Chytrý et al. 2009). An additional level of complexity is added in studies dealing with grid data where both CWM and internal sample attributes (e.g. species richness derived from community data) are spatially autocorrelated due to the spatial coherence of species distribution (Hawkins et al. 2017).

### Category B – species attributes linked to species composition

Studies in this category explicitly assume that the species attributes are linked to species composition, and the test focuses on the link of sample attributes to species composition. The null hypothesis states that sample attributes are not linked to species composition, while alternative hypothesis states they are. Examples are trait-based studies asking whether species traits can explain the effect of environmental filtering on species abundance in a community. These studies operate with an assumption that traits (species attributes) are functional, i.e. they influence the abundance of species in a community, and the question which is tested is whether the sample attributes (environmental variables) act as an environmental filter on species abundance. Studies using internal species attributes (derived from species composition, e.g. as the weighted-mean of sample attributes or as scores on ordination axes) also belong to this category.

### Category C – no assumption about the link between species or sample attributes and species composition

This category includes mostly observational studies without prior assumptions or expectations about a link between any of the matrices. The null hypothesis states that there is no link between species or sample attributes and the matrix species composition. To reject this null hypothesis means to prove that both species and sample attributes are linked to species composition. Empirical studies describing the general relationship between sample attributes and species attributes, without explicitly or implicitly acknowledging some underlying assumption or mechanism, belong to this category. Examples are studies relating the CWM of functional traits to environmental variables without a clear assumption that traits are functional, allowing to question whether particular traits are linked to species composition or not. Studies with species indicator values relating mean indicator values to environmental variables also fit this category (e.g. answering the question of whether Ellenberg indicator values for soil reaction *per se* are good predictors of measured soil pH).

## Illustration of the bias and its dependence on beta diversity

If we test hypotheses from categories A and C by the weighted-mean approach with the standard test, results may be highly biased, both regarding the estimated model parameters and the inflated Type I error rate (Zelený & Schaffers 2012; Peres-Neto et al. 2016; Hawkins et al. 2017). In the next section, I will illustrate this bias using simulated community data, where each community data set will be accompanied with (vectors of) species and sample attributes linked (or not) to species composition. To show also the dependence of the bias on the heterogeneity of species composition, I will generate sets of species composition matrices of increasing beta diversity. Later, I will use the same simulated data sets to demonstrate the performance of available statistical solutions.

### Description of 2D simulated community data set

The algorithm generating community data is an extension of the one proposed by Fridley et al. (2007) and is structured by two virtual ecological gradients. I call it *2D simulated community data set* throughout this paper, to distinguish it from an alternative algorithm introduced by Dray & Legendre (2008) structured by only one virtual ecological gradient (and called *1D simulated community data set* here). Along each virtual gradient, a certain number of unimodal species response curves were generated, where response curve quantifies the potential probability that individual found in the certain location of the gradient would be assigned to given species. Community samples were then generated by randomly selecting locations along the gradients and assigning given the number of individuals into species according to species probabilities at given gradient location. Sample locations along the first gradient were used as sample attributes, while optima of species response curves along the first gradients were used as species attributes. The function of the second gradient is to modify the beta diversity of the dataset; increasing the length of the second gradient (along to the increasing number of species) results in increased beta diversity of the species composition matrix (Appendix S1: Table S1 and Fig. S1). The range of species niche widths was between 500 and 1000 units, and niche widths were generated independently for each gradient. The length of the first gradient was arbitrarily set to 1000 units, while the length of the second gradient varied between 1000 to 10 000 units. I assumed that 1000 units of the second gradient represent one community, i.e. enlarging the second gradient from 1000 to 10 000 units (by steps of 1000 units) generates a set of data sets with 1 to 10 communities. For more details, see Appendix S1.

As a result, each simulated community data set includes a matrix of sample attributes (**R**), species composition (**L**) and species attributes (**Q**), in which sample attributes and species attributes are linked to species composition. In the next step, the link between species or sample attributes and species composition (or both) was broken by the permutation of attributes to create four scenarios (Fig. 1, identical with scenarios 1–4 of Dray & Legendre 2008): 1) both sample and species attributes are linked to species composition; 2) sample attributes are linked to species composition, species attributes are not; 3) species attributes are linked to species composition, sample attributes are not; 4) none of the species or sample attributes is linked to species composition. For studies from category A defined above, scenario 2 represents the null hypothesis, for category B scenario 3 is the null hypothesis, and for category C the scenarios 2, 3 and 4 represent alternative states of the null hypothesis (Table 1 and Fig. 1). Scenario 1 represents the power test for all three categories (i.e. it measures the probability of getting significant results if the alternative hypothesis is true). Note that in the simulated data example, sample and species attributes are matrices with a single column (vectors), yet for simplicity, I keep using the matrix notation, i.e. **R** and **Q** instead of **r** and **q**.

For the comparison with results of the study by Peres-Neto et al. (2016), I calculated all analyses also using *1D simulated community data set* (results available in Appendix S3). This model generated rather homogeneous communities (Appendix S1: Table S1 vs. Appendix S3: Table S2), and the increase in beta diversity was achieved by narrowing the niche breadth of individual species (keeping the gamma diversity of the data set constant but decreasing the mean alpha diversity). In contrast, in 2D simulated community data set the increase in beta diversity was achieved by prolonging the second virtual gradient (which increases gamma diversity while keeping the mean alpha diversity rather constant).

All analyses were conducted using R-project (version 3.3.3, R Foundation for Statistical Computing, Vienna, Austria, https://www.R-project.org/); complete R-script is available in Appendix S4, and all functions are in R-package *weimea* (abbreviation for the *weighted mean*; source code of version 0.64 is in Appendix S5, actual version can be found at https://github.com/zdealveindy/weimea).

### Weighted-mean approach with standard test applied on simulated data

For each of the four scenarios (1–4) I created ten levels of beta diversity, and for each combination of *scenario* × *level of beta diversity*, I created 1000 datasets (4 scenarios × 10 levels of beta diversity × 1000 replications = 40 000 data sets). For each data set, I calculated the CWM of species attributes, related it to sample attributes using Pearson’s *r* correlation and tested its significance using the parametric *t*-test (for additional results for least-square regression and *r*^*2*^ see Appendix S2: Fig. S1). For each level of community beta diversity in each scenario, I counted the proportion of correlations significant at *α* = 0.05 (note that this proportion is identical to the proportion of significant regressions).

From the three scenarios with no direct link between species and sample attributes (scenarios 2, 3 and 4), analysis of data generated by scenario 2 reveals the bias – the correlation coefficient deviates from zero more than in other cases (Fig. 2), and the test of significance shows an inflated Type I error rate (Fig. 3). This bias decreases with increasing beta diversity of the species composition matrix (Fig. 2 & 3, Scenario 2): for the most homogeneous data set (*level of beta diversity* = 1), the range of Pearson’s *r* correlation coefficients (expressed as 2.5% and 97.5% quantiles) is between -0.761 and 0.782, with 58% of correlations significant, while for the most heterogeneous data set with a high beta diversity (*level of beta diversity* = 10) the range of Pearson’s *r* values is between -0.399 and 0.390 (2.5 and 97.5% quantile range), with 17% of correlations significant. For comparison, the most homogeneous datasets in scenarios 3 and 4 have the values of *r* on average between -0.286 and 0.275 (2.5 and 97.5% quantile range) with the number of significant results 5.7%, i.e. close to expected 5%. Similarly inflated are the values of coefficient of determination (*r*^*2*^; Appendix S2: Fig. S1, Scenario 2) calculated by least-square linear regression. Applying the standard test on the simulated community data set of Dray & Legendre (2008) shows analogously biased results (Appendix S3: Table S1 and Fig. S2a).

**Figure 2.**
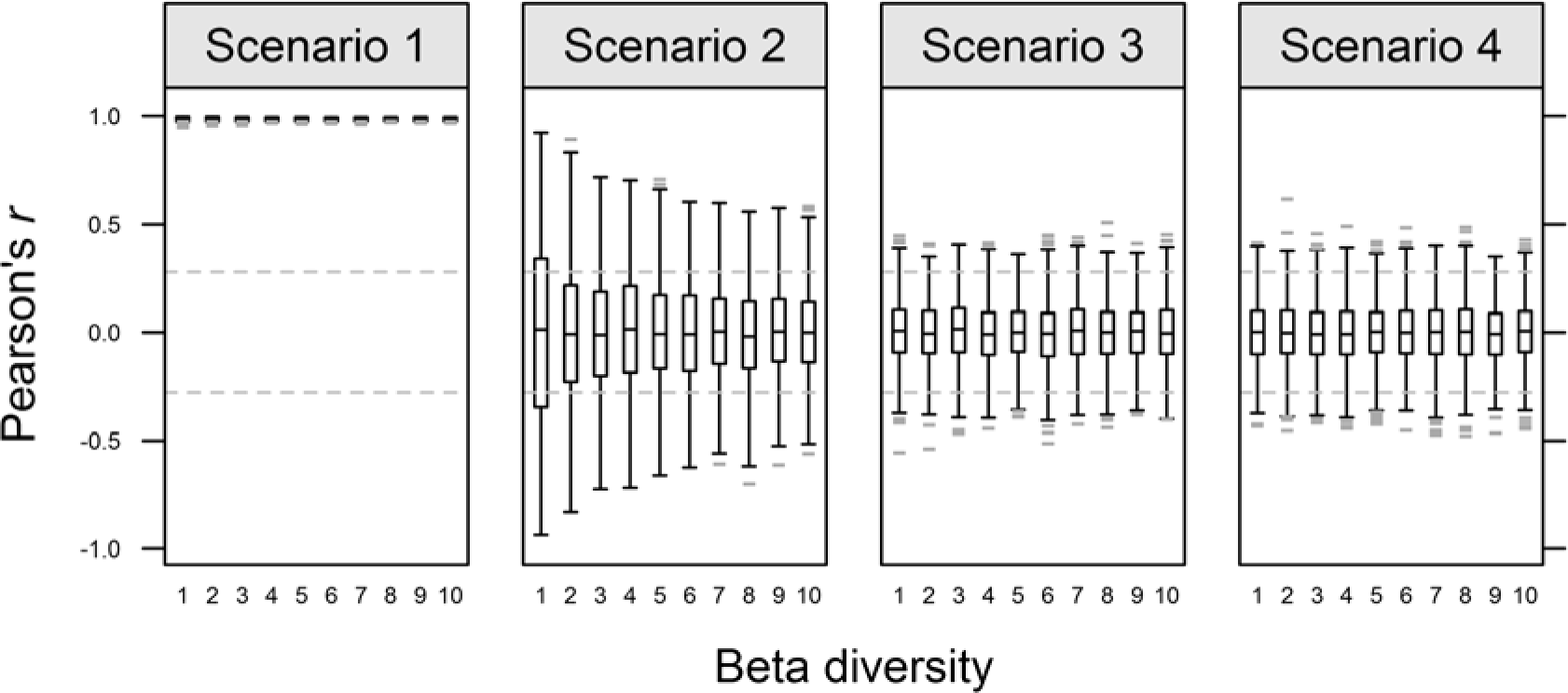
Pearson’s *r* correlation coefficients among CWM and sample attributes for each of the four scenarios and ten levels of beta diversity (1000 correlations for each combination have been conducted). Grey horizontal bars are outliers.

**Figure 3.**
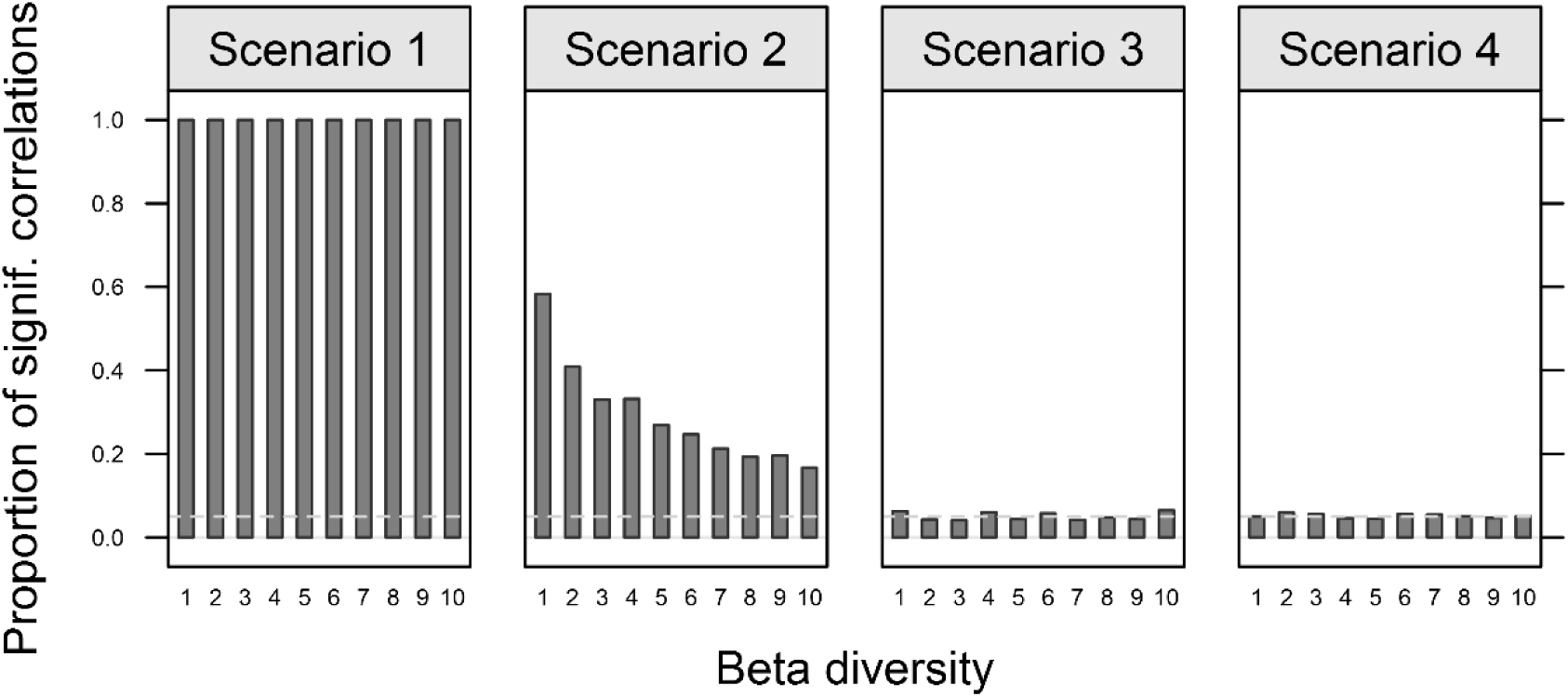
The proportion of significant correlations (P < 0.05) between CWM and sample attributes, tested by standard parametric *t*-test. For each of the four scenarios and ten levels of beta diversity, 1000 simulated community data sets have been generated and tested.

## Available solutions and their comparison

To my knowledge, there are two approaches that have been used to solve the bias in the weighted-mean approach. One is the *modified permutation test*, which was introduced by Zelený & Schaffers (2012) in the context of relating the CWM of Ellenberg-like species indicator values with internal variables (e.g. ordination scores or species richness calculated from the same species composition data set). Another one is the *sequential permutation test* using the *fourth-corner statistic*, introduced first in the electronic appendix of the study by Peres-Neto et al. (2012, Appendix A) and later in a re-elaborated version in Peres-Neto et al. (2016). Here I review the strengths and weaknesses of both approaches, test their performance using simulated community data and suggest guidelines for their use. Both 2D and 1D simulated community data sets have been used, with results of the two-gradient version reported in the main paper and those of one-gradient version in Appendix S3. Note that studies in category B are not prone to the bias if the weighted-mean approach with the standard test is used, and reviewed solutions are therefore relevant only for studies in categories A and C.

### Modified permutation test: comparison with the results of a null model

Standard permutation test of the relationship between sample attributes and CWM of species attributes compares observed test statistic (e.g. t-value for correlation or F-value for regression) with expected null distribution of this test statistic generated by repeatedly randomizing one of the compared variables (e.g. by randomizing CWM values between samples). *Modified permutation test* modifies the way how the null distribution is generated, and instead of generating CWM values between samples, it calculates CWM on species attributes randomized among species (or randomly generated ones). CWM calculated from randomized species attributes (CWM_rand_) inherit the same level of compositional autocorrelation as CWM calculated from the real species attributes (CWM_obs_) because they are calculated by the same algorithm from the same species composition matrix. This is analogous to testing the relationship between spatially autocorrelated variables using toroidal shift, when one spatially explicit variable is permuted in a way that it preserves the original degree of spatial autocorrelation (Fortin & Dale 2005), or, alternatively, random variables with the same degree of spatial autocorrelation as that of the original one can be generated (Deblauwe et al. 2012).

I used 2D and 1D simulated community data sets to calculate the correlation of CWM sample attributes for all four scenarios in communities of increasing beta diversity and tested the significance of this correlation using the modified permutation test. Results on 2D data set show that in contrast to the standard permutation test (Fig. 3), the originally inflated Type I error rate in the case of scenario 2 disappears (Fig. 4). At the same time, in the case of scenario 3 the test is slightly conservative for data sets of low beta diversity. The same conclusion applies if the modified permutation test is used on 1D simulated community data set, in which the results for scenario 3 are even more conservative (almost no significant correlations, Appendix S3: Table S1 and Fig. S2a), since the community data set has a rather low beta diversity (Appendix S3: Table S2). Additional detail power analysis on the 1D simulated community data set with added random noise reveals that the modified permutation test loses power when both sample size and species number decrease (Appendix S3: Fig. S1a), and also when species tolerances increase (Appendix S3: Fig. S1b; note that with increasing species tolerance, beta diversity of the species composition matrix decreases).

**Figure 4.**
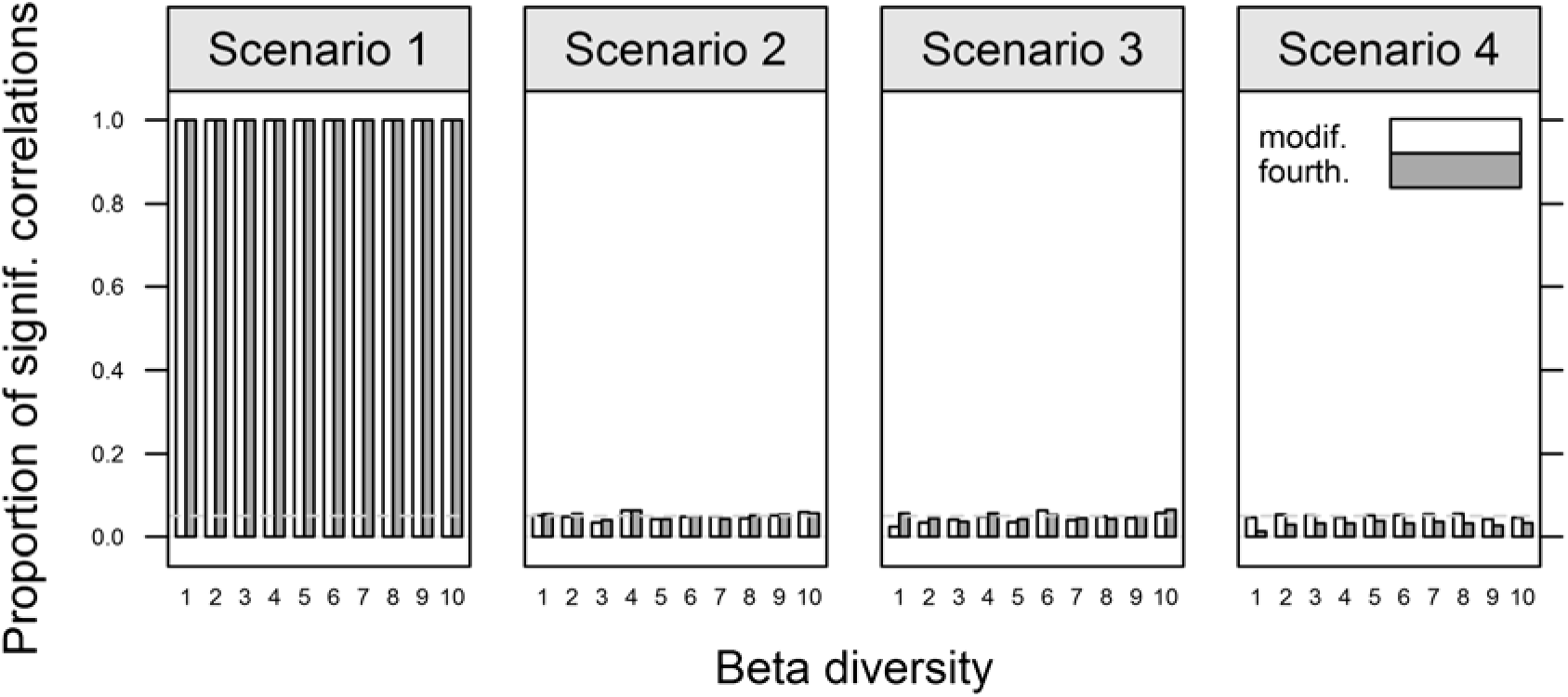
The proportion of significant correlations (P < 0.05) between CWM and sample attributes, tested by modified permutation test (white bars) and sequential test with fourth-corner *r* statistic (gray bars). For each of the four scenarios and ten levels of beta diversity, 1000 simulated community data sets have been generated and tested by each method.

The modified permutation test is suitable for testing hypotheses in both categories A and C, since for both scenario 2 is relevant for testing the null hypothesis. In the case of category C, however, it is not suitable for data sets with very low beta diversity, for which results of modified permutation test in scenario 3 are overly conservative.

### Sequential permutation test with the fourth-corner statistic

Dray & Legendre (2008) noted that the fourth-corner statistic *r*, introduced by Legendre et al. (1997), is “equal to the slope of the linear model, weighted by total species abundance, with the niche centroids as the response variable and the species trait as the explanatory variable.” This analogy was further elaborated by Peres-Neto et al. (2012, Appendix A) and Peres-Neto et al. (2016), who presented an algorithm for how to use the *fourth-corner* statistic *r* in the weighted-mean approach. In short, both **R** and **Q** matrices are first centered by the weighted mean of row sums of **L** (in the case of **R**) and column sums of **L** (in the case of **Q**), and then standardized. The fourth-corner *r* statistic is then the slope of weighted regression between the weighted mean of centered plus standardized **Q** and centered plus standardized **R**, weighted by row sums of **L**. The main advantage of the fourth-corner statistic is the option to use the *sequential permutation test* introduced by ter Braak et al. (2012), which combines results of tests based on permuting sample attributes (model 2 in Legendre et al. 1997) and species attributes (model 4). If the first test is significant, then the second test is done, and overall significance of the result is equal to the higher of these two tests’ *P*-values. When applied to the 2D simulated community data set, this test gives unbiased results for all scenarios (Appendix S2: Fig. S3), although being more conservative in the case of homogeneous data sets in scenario 4, which is relevant for category C (results calculated on the 1D simulated data set also confirm this finding, see Appendix S3:

Table S1 and Fig. S2b). Power analysis (Appendix S3: Fig. S1c,d) reveals a performance very similar to that of the modified permutation test. The sequential test with the fourth-corner statistic is, therefore, suitable for testing hypotheses from all three categories, although in the case of category B it is not needed (standard permutation test gives unbiased results) and in the case of category C it is overly conservative for homogeneous community data sets (scenario 4 on Fig. 4). A disadvantage is that the sequential test with the fourth-corner statistic is restricted only to the weighted regression/correlation between centered and standardized species and sample attributes, weighted by row sums of a species composition matrix (**L**). This may not be appropriate e.g. in the case of presence-absence species composition data if CWM are related to species richness (as sample attributes). The regression relating CWM to species richness would then be weighted by species richness (row sums in presence-absence species composition data equal to species richness), in which case more species-rich samples would have a higher weight in the analysis (Hawkins et al. 2017). Sequential test with the fourth-corner statistic is, therefore, more like a special case of weighted-mean approach, which also includes other methods such as non-weighted regression, correlation or ANOVA, does not require standardizing species and sample attributes and does not weight the samples by sums of their species abundances.

## Discussion

The main motivation of this study was to show that the results of the weighted-mean approach critically depend on the correct decision regarding the test used for statistical inference. To help in this decision process, I suggested that each ecological question analyzed by weighted-mean approach should be classified into one of the three categories, given the explicit (or implicit) assumptions about the role of species and sample attributes. For each category, I suggested an optimal strategy for testing the significance of the relationship between the CWM and sample attributes, summarized in Table 1. The choice of the appropriate category is not always straightforward. For example, trait studies testing whether an environment is filtering the species into a community via their functional traits routinely assume that such traits are functional, and in the weighted-mean approach they are considered as linked to species composition (category B). However, this assumption may not always be justified; included traits are often those readily available in databases and/or those which are relatively easy to measure, but these do not necessarily need to be the functional ones (Mlambo 2014). In the case of compositionally relatively homogeneous data sets, even the traits with no ecological meaning may show a high and significant relationship to environmental variables if tested by standard tests. I believe that this calls for a revision of such commonly applied practice.

The analogy between the bias in the weighted-mean approach to the bias in the analysis of spatially autocorrelated variables suggests some other alternatives to reduce or remove the bias. One is to stratify the data set to reduce redundancy in species composition among samples and increase the overall beta diversity of the compositional data set, e.g. by removing one sample from pairs of samples with similar species composition. Although methods for stratification based on species composition are available (e.g. Lengyel et al. 2011), this potentially results in throwing out a large number of expensive data. Another option would be to apply some correction for effective degrees of freedom in analysis, analogous to a method estimating the effective number of samples in the case of autocorrelated variables (Dutilleul 1993), or to apply methods capable of dealing with autocorrelated residuals (analogy of geographically weighted regressions).

The power test using the simulated data set showed that the power of both the modified permutation test and the sequential permutation test with the fourth-corner statistic decreases with a decrease in the number of species and/or the number of samples. This makes these tests less suitable for smaller and relatively homogeneous data sets with few species (e.g. less than 40) since the probability of Type II error (i.e. not rejecting the null hypothesis, which is false) strongly increases. Additionally, in the case of compositional data sets with low beta diversity, the modified permutation test is overly conservative for scenario 3, while the sequential permutation test with the fourth-corner statistic is conservative for scenario 4. Both tests are therefore less suitable for testing hypotheses in category C in the case that compositional data set is rather homogeneous.

In this study, I explicitly ignored intraspecific variation in species attributes, focusing only on the use of data set-wide mean species attribute values. Indeed, intraspecific variation may be important, e.g. in the context of functional traits, where the intraspecific variation gains increasing attention (Albert et al. 2012). The question whether the inclusion of intraspecific variation (e.g. by including trait values that are sample-specific, not data set-wide) influences the potential bias reported in this study requires further examination and goes beyond this study. I assume, however, that including another source of variation (species-level variation in species attributes) does not remove the problem of the bias itself but makes the estimation of the bias and its correction more complex.

Finally, the relevant consideration is whether the weighted-mean approach is actually the best analytical solution for the question we aim to answer. In some cases, the question is explicitly focused on relating community-level values of species attributes, like mean Ellenberg-like species indicator values (serving as an estimate of ecological conditions for individual sites) or the CWM of traits (as one of the functional-diversity metrics and as a community-level trait value), and the use of the weighted-mean approach is fully justified. In other cases, when the question is focused on relating individual species-attributes to sample attributes, the weighted-mean approach may not be the best analytical choice. A better solution may be to use alternative options, such as the fourth-corner (Legendre et al. 1997) or RLQ (Dolédec et al. 1996) analysis, for which the problem of inflated Type I error rate and choice of the suitable permutation test have already been solved.

## Conclusions

In this study, I draw attention to the problem of the weighted-mean approach, which I believe is largely overlooked and not acknowledged, although it represents a source of potentially serious misinterpretations. Since in certain fields of vegetation ecology the weighted-mean approach is gaining increasing momentum (e.g. in functional ecology with the CWM of species functional traits as one of the functional-diversity indices), I suggest that the time is ripe to critically assess in which situations and for which types of hypotheses the commonly used standard parametric or permutation tests are inappropriate, since they yield results that may be overly optimistic.

## Acknowledgements

This study was supported by the Czech Science Foundation (P505/12/1022) and Ministry of Science and Technology, Taiwan (MOST 105-2621-B-002-004). My thanks go to Bill Shipley, Cajo ter Braak and several anonymous reviewers for critical comments on previous versions of this manuscript, which motivated me to several times heavily rework it. Thanks also go to Pedro Péres-Neto and Stéphane Dray for (emotional) discussion of differences between my modified permutation test solution and their fourth-corner one during the ISEC 2014 conference in Montpellier.

## Supporting information

**Appendix S1.** Description of an algorithm generating simulated community data along two environmental gradients (*2D simulated community data set*).

**Appendix S2.** Weighted-mean approach applied to 2D simulated community data sets: additional results.

**Appendix S3.** Evaluation of permutation tests using 1D simulated community data set from Dray & Legendre (2008).

**Appendix S4.** R-code for all analyses.

**Appendix S5.** Source code for the R library *weimea*, version v. 0.64 (actual version can be found on https://github.com/zdealveindy/weimea/).

